# SARS-CoV-2 causes human BBB injury and neuroinflammation indirectly in a linked organ chip platform

**DOI:** 10.1101/2021.10.05.463205

**Authors:** Peng Wang, Lin Jin, Min Zhang, Yunsong Wu, Zilei Duan, Wenwen Chen, Chaoming Wang, Zhiyi Liao, Jianbao Han, Yingqi Guo, Yaqiong Guo, Yaqing Wang, Ren Lai, Jianhua Qin

## Abstract

COVID-19 is a multi-system disease affecting many organs outside of the lungs, and patients generally develop varying degrees of neurological symptoms. Whereas, the pathogenesis underlying these neurological manifestations remains elusive. Although *in vitro* models and animal models are widely used in studies of SARS-CoV-2 infection, human organ models that can reflect the pathological alterations in a multi-organ context are still lacking. In this study, we propose a new strategy to probe the effects of SARS-CoV-2 on human brains in a linked alveolus-BBB organ chip platform. The new multi-organ platform allows to recapitulate the essential features of human alveolar-capillary barrier and blood-brain barrier in a microfluidic condition by co-culturing the organ-specific cells. The results reveal direct SARS-CoV-2 exposure has no obvious effects on BBB chip alone. While, infusion of endothelial medium from infected alveolus chips can cause BBB dysfunction and neuroinflammation on the linked chip platform, including brain endothelium disruption, glial cell activation and inflammatory cytokines release. These new findings suggest that SARS-CoV-2 could induce neuropathological alterations, which might not result from direct viral infection through hematogenous route, but rather likely from systemic inflammation following lung infection. This work provides a new strategy to study the virus-host interaction and neuropathology at an organ-organ context, which is not easily obtained by other *in vitro* models. This will facilitate to understand the neurological pathogenesis in SARS-CoV-2 and accelerate the development of new therapeutics.

**SUMMARY:**

1. A linked human alveolus-BBB chip platform is established to explore the influences of SARS-CoV-2 on human brains in an organ-organ context.
2. SARS-CoV-2 infection could induce BBB injury and neuroinflammation.
3. The neuropathological changes are caused by SARS-CoV-2 indirectly, which might be mediated by systemic inflammation following lung infection, but probably not by direct viral neuroinvasion.

## INTRODUCTION

The coronavirus disease 2019 (COVID-19) is a multi-organ disease. Although the symptoms of COVID-19 are predominantly concentrated in human respiratory system, obvious organs dysfunction outside of the lungs are detected, including intestine, heart, kidney, liver and brain ^1-4^. Of note, accumulating clinical evidences showed 30-40% COVID-19 patients develop obvious neurological symptoms, such as headache, dizziness, psychosis, hypogeusia, hyposmia, cerebrovascular injury, seizures and encephalitis ^5-8^. In human brain, there is a highly selective blood-brain barrier (BBB), which controls the transport of nutrients and metabolites between the systemic circulation and the nervous system ^9,10^. Also, BBB functions as a protective barrier to prevent toxins and pathogens (including virus) from accessing the brain parenchyma. In some brain infectious diseases, neurotropic viruses can infect and disrupt BBB, and further invade human brain through the hematogenous route, such as HSV-1 and Zika virus ^11-13^. However, whether SARS-CoV-2 can cross BBB to invade brain through the similar route remains elusive.

In the past year, great efforts have been made to investigate the influences of SARS-CoV-2 on brains. By examining autopsy samples from COVID-19 patients, several pathological studies revealed cerebral injuries in patients’ brains, such as cerebrovascular damage and glial activation ^14,15^. Some studies using animal models, reported SARS-CoV-2 can be detected in brains after intranasal administration of virus, indicating potential neuroinvasion of SARS-CoV-2 ^16-18^. As a new type of *in vitro* model, brain organoids contain multiple cell types and have been widely used in studies of the SARS-CoV-2 neurotropism, showing different susceptibility of various brain cells for the viral infection, including neurons, neural progenitor cells and astrocytes ^19-24^. However, some limitations constrain interpretations of these studies. As for autopsy brain tissues, limited availability of samples from COVID-19 patients and variability in patient presentation restricts their large-scale application ^14,15^. Classic animal models, such as rodents, are different from humans in physiological structure, and are limited in accurately recapitulating neuropathological changes of COVID-19 in humans ^25,26^. Regarding brain organoids, functional brain barriers and immune cells are absent, which makes it difficult to accurately recapitulate viral exposure and host immune responses in a physiological state. Especially, given that COVID-19 is a multi-organ disease, and damaged organs are closely associated during disease pathogenesis. Human organ models that can reflect pathological changes at systematic level is still lacking.

Microfluidic organ chip models, emerging as *in vitro* microphysiological systems, possess the potential to recreate functional units from the native human organs or tissues. It can resemble the organ physiology in a human-relevant manner by mimicking the *in vivo*-like cellular microenvironment factors, such as cell-cell interactions and mechanical fluid flow. This technology has been developed to create a variety of organ models, such as the lung, liver, heart and BBB ^10,27-33^, holding great potential in disease modeling and drug testing. More recently, several organ chips have been leveraged to study host responses to SARS-CoV-2 infection and assess the antiviral effect of candidate drugs ^34-39^.

In this study, we proposed a new strategy to create a linked alveolus-BBB chip platform that allows to explore the effects of SARS-CoV-2 infection on human BBB in a multi-organ context. The multi-organ chip platform was created by linking an alveolus chip and BBB chip via fluidic control. Human peripheral blood immune cells were introduced into the multi-organ platform to simulate the cytokine storm following SARS-CoV-2-induced lung infection. Our study showed, direct SARS-CoV-2 exposure had no visible effects on BBB chip alone. While, SARS-CoV-2 induced BBB injury and inflammation indirectly following the systematic inflammation caused by lung infection on the linked chip platform. Collectively, our findings provide a possible explanation for pathogenesis underlying SARS-CoV-2-induced neurological symptoms, which does not result from direct viral infection through hematogenous route, but rather likely from systemic inflammation following lung infection.

## RESULTS

### Establishing a linked human alveolus-BBB chip platform for studies of viral infection

COVID-19 is a multi-organ disease. Although lung is the major target organ for SARS-CoV-2, obvious neurological symptoms can be diagnosed in a large population of COVID-19 patients (Figure 1A). In order to probe the effects of SARS-CoV-2 infection on human brains in an organ-organ context, we plan to establish a linked human alveolus-BBB organ chip model system that enables to recapitulate the essential features of human alveolar-capillary barrier and blood-brain barrier in a physiologically relevant manner (Figure 1B).

**Figure 1.**
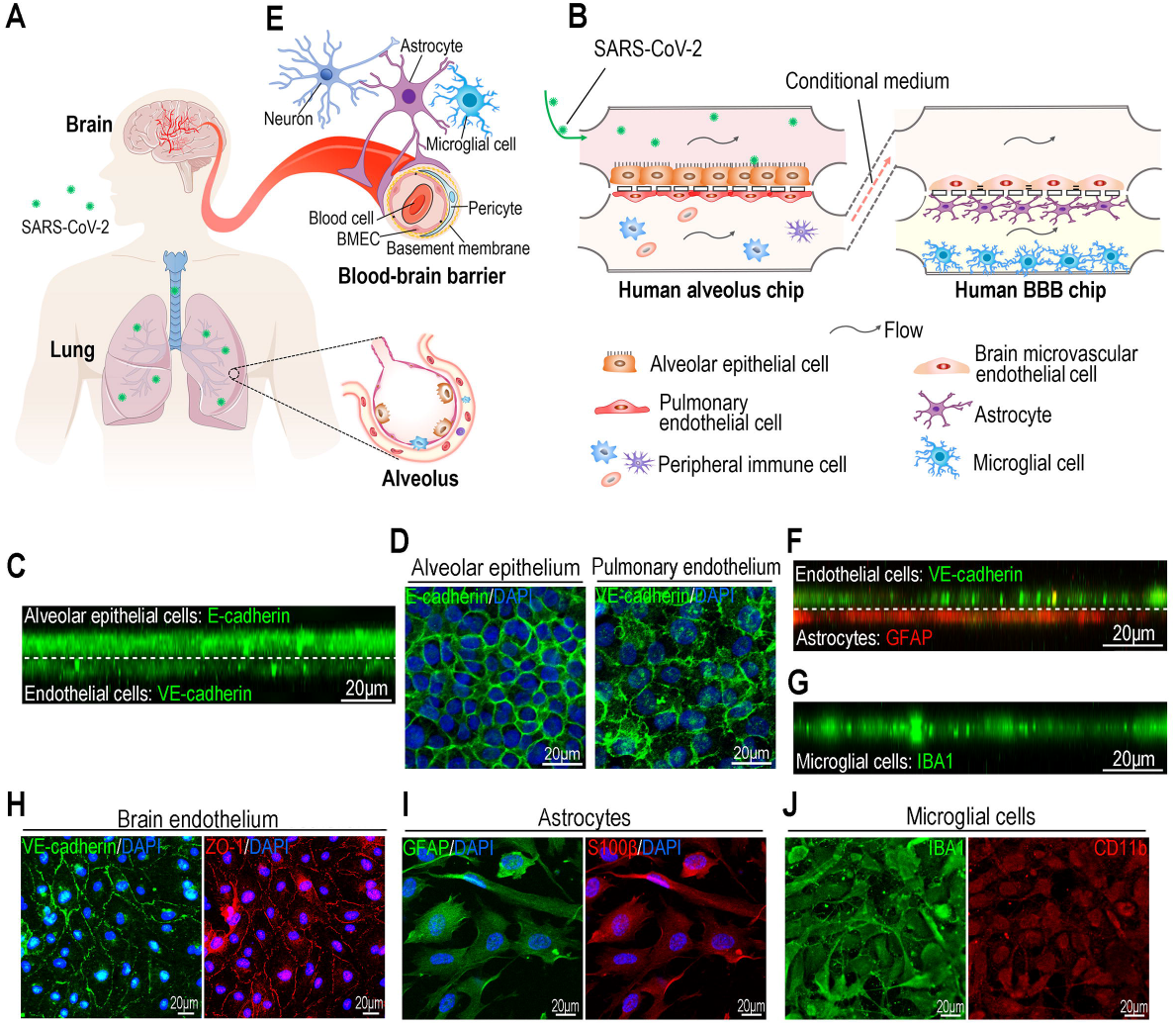
A linked alveolus-BBB organ chip platform for studies of viral infection. (A) Schematic diagram of human body exposed to SARS-CoV-2. (B) Schematic description of a linked alveolus-BBB chip platform, consisting of a human alveolus chip and a human BBB chip. (C) A side view of confocal immunofluorescent image showing alveolar-capillary barrier interface on chip device. The alveolar-capillary barrier interface was formed by co-culture of human alveolar epithelial cells (HPAEpiC; E-cadherin staining) and human pulmonary microvascular endothelial cells (HULEC-5a; VE-cadherin staining) on the opposite sides of porous membrane under fluid flow conditions. (D) Confocal immunofluorescent micrographs showing the alveolar epithelium (E-cadherin) and pulmonary endothelium (VE-cadherin) cultured on chip device for 3 days. (E) Illustration of human BBB comprising brain microvascular endothelial cells (BMECs), pericytes and perivascular astrocytes. (F) A side view of confocal immunofluorescent image showing blood-brain barrier interface on BBB chip device. The BBB interface was formed by co-culture of brain microvascular endothelial cells (HBMEC; VE-cadherin staining) and astrocytes (HA; GFAP staining) on the opposite sides of porous membrane under fluid flow conditions. (G) A side view of confocal immunofluorescent image showing microglia (HMC3; IBA1 staining) cultured on the bottom side of CNS channel. (H) Confocal immunofluorescent micrographs showing the human brain microvascular endothelial cells cultured on BBB chip for 3 days immunostained for VE-cadherin and ZO-1. (I) Confocal immunofluorescent micrographs showing the human astrocytes cultured on BBB chip for 3 days immunostained for GFAP and S100β. (J) Confocal immunofluorescent micrographs showing the human microglia cultured on BBB chip for 3 days immunostained for IBA1 and CD11b.

The human alveolus chip consisted of an upper epithelial channel and a lower vascular channel separated by a porous polyethylene terephthalate (PET) membrane (10 μm thick, 2 μm diameter). The alveolar-capillary barrier interface was established by co-culturing human alveolar epithelial cells (HPAEpiC) and human pulmonary microvascular endothelial cells (HULEC-5a) on the porous membrane (precoated with fibronectin) under microfluidic condition (100 μL/h) (Figure 1C, D) ^38^.

In human brain, brain microvascular endothelial cells (BMECs) are the primary anatomical component of the BBB and line the brain capillaries. They work in concert with basement membrane, perivascular astrocytes and pericytes, as well as neighboring neurons and microglia to form the neurovascular unit (Figure 1E) ^10,40-42^. To reconstitute the human BBB *in vitro*, we fabricated a microfluidic BBB chip, which consisted of an upper vascular channel and a parallel CNS channel separated by the porous PET membrane. Human brain microvascular endothelial cells (HBMEC) and human astrocytes (HA) were seeded on the opposite sides of porous membrane, mimicking the BBB interface *in vivo*, and the human microglial cells (HMC3) were cultured on the lower side of CNS channel. Both channels were perfused at a flow velocity of 100 μL/h for 3 days. Side view of confocal microscopic images revealed an intact tissue interface was formed on the porous membrane identified by VE-cadherin (adherens junction protein) in brain endothelial cells and GFAP (astrocytic marker) in astrocytes (Figure 1F), and a microglial layer (IBA1) was formed on the bottom side of CNS channel (Figure 1G). In brain endothelium, well developed adherens junctions (VE-cadherin) and tight junctions (ZO-1) were formed (Figure 1H). As for glial cells, obvious astrocyte markers (GFAP, S100β) and microglial markers (IBA1, CD11b) were detected on the astrocytic layer (Figure 1I) and microglial layer (Figure 1J), respectively. These results reveal the integrity of BBB interface was formed under the microfluidic condition.

### Direct exposure of SARS-CoV-2 showed no visible effects on BBB chip alone

In order to evaluate the viral invasiveness to human brain via infecting BBB, we conducted direct SARS-CoV-2 exposure test on individual BBB chip. The individual alveolus chip was infected with SARS-CoV-2 (MOI=1) in parallel as a positive control. For SARS-CoV-2 exposure test on BBB chip, we chose a MOI (MOI=1) that is much higher than physiological amount of SARS-CoV-2 exposure in peripheral blood for human brains. Initially, the expression of ACE2 and TMPRSS2 in cells which SARS-CoV-2 utilizes for cellular entry, was determined by Western blot. It appeared both proteins were expressed most highly in HPAEpiC cells (human alveolar epithelial type II cells-derived cell line), followed by two vascular endothelial cell lines (HULEC-5a, HBMEC) and glial cell lines (HA, HMC3) (Figure 2A). This was consistent with scRNA-seq analysis that ACE2 and TMPRSS2 are specifically highly co-expressed in human alveolar epithelial type II cells ^43,44^.

**Figure 2.**
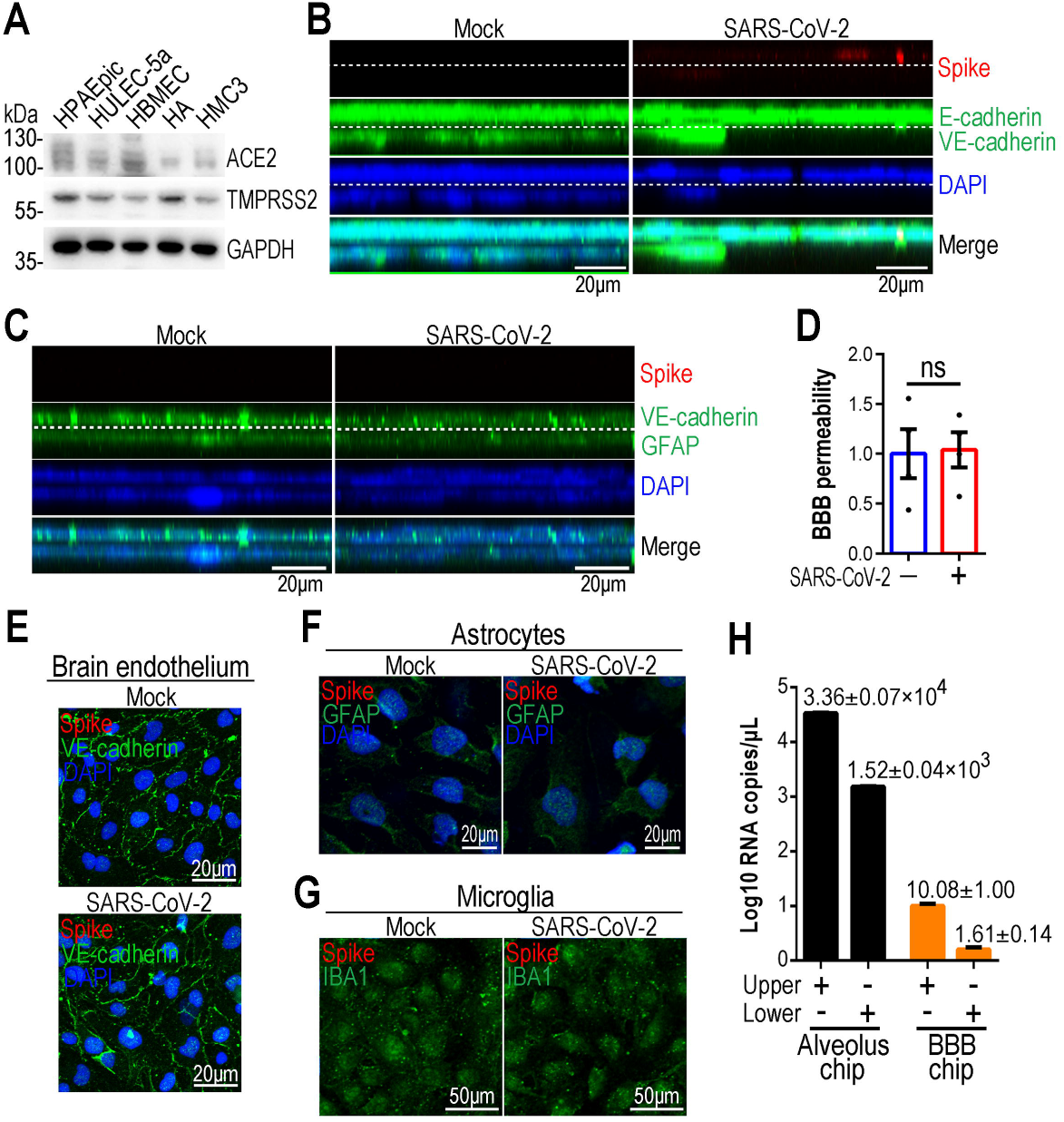
Testing direct exposure of SARS-CoV-2 on the BBB chip alone. (A) Western blot showed the protein levels of ACE2 and TMPRSS2 in five cell lines. The result was a representative blot from three independent experiments. GAPDH was used as a loading control. (B) Representative side views of the alveolar-capillary barrier interface 3 days post SARS-CoV-2 infection in the individual alveolus chip. SARS-CoV-2 infection and replication was identified primarily in alveolar epithelium (E-cadherin; green) by SARS-CoV-2 Spike protein expression (red) (n=3). (C) Representative side views of the blood-brain barrier interface 3 days post SARS-CoV-2 infection in the individual BBB chip model. No SARS-CoV-2 infection or replication was identified in both endothelial layer (VE-cadherin; green) and astroglial layer (GFAP; green) by SARS-CoV-2 Spike protein expression (red) (n=4). (D) FITC-dextran permeability assays showed the effect of direct SARS-CoV-2 infection on BBB integrity (n=4). The FTIC-dextran concentration of lower channel in mock-infected group was normalized to 1, and that of SARS-CoV-2 group was presented as the multiples of FTIC-dextran concentration in the control group. Data are presented as mean±SEM and analyzed by unpaired Student’s t test. (E) Representative confocal micrographs of brain endothelial cells immunostained for VE-cadherin and Spike following SARS-CoV-2 infection on BBB chip (n=4). (F) Representative confocal micrographs of astrocytes immunostained for GFAP and Spike following SARS-CoV-2 infection on BBB chip (n=4). (G) Representative confocal micrographs of microglia immunostained for IBA1 and Spike following SARS-CoV-2 infection on BBB chip (n=4). (H) qRT-PCR results showing the viral loads for culture supernatants of human alveolus chip and BBB chip. The number on each bar indicates the viral RNA copies per volume of culture. Data represent three independent experiments and were presented as mean±SEM.

Three days post-infection, obvious infection and replication of SARS-CoV-2 was detected on the individual alveolus chip identified by viral Spike protein (Figure 2B), which was consistent with our previous study ^38^. However, no obvious SARS-CoV-2 infection was detected on the individual BBB chip (Figure 2C). No visible changes in BBB permeability were observed following direct viral exposure either (Figure 2D). In brain endothelial cells, adherent junctions visualized by VE-cadherin were well preserved in the infected chip (Figure 2E). In the lower CNS channel of the infected chip, no viral infection and cell morphological changes were observed in both astrocytes and microglia as well (Figure 2F, G).

To exclude the possibility that SARS-CoV-2 replication in BBB is very low and below the limits of detection by immunofluorescence, culture mediums of infected BBB chips were analyzed by qRT-PCR for viral RNA detection, which is a more sensitive method. The result showed (Figure 2H), viral loads of infected BBB chips were much lower than those of infected alveolus chips, indicating BBB is less permissive to SARS-CoV-2 direct infection, and the probability of direct neuroinvasion via BBB is low.

### SARS-CoV-2 caused BBB injury indirectly following lung infection on the linked chip platform

As direct viral exposure of SARS-CoV-2 had no visible effects on BBB chip alone, we then assumed neuropathological changes may be caused by the systematic inflammation following lung infection in an indirect manner.

To verify the hypothesis, we initially collected endothelial medium from the mock- or SARS-CoV-2 infected human alveolus chips (MOI=1), and then utilized it as conditional medium to treat BBB chips. Here, we chose influenza H1N1 virus (PR8 strain), as a reference in parallel (MOI=1) (Supplementary Figure 1A). To test whether SARS-CoV-2 infection can induce a cytokine storm that commonly happens in severe COVID-19 cases, cytokines in culture supernatants of alveolus chips were detected 4 days after infection. The results showed 10 cytokines (including IFN-α2, IL-2, IFN-γ, IL-7, IL-1RA, IL-8, TNF-α, CXCL10, CCL3 and IL-10) were significantly released in the epithelial channels of SARS-CoV-2-infected alveolus chip (Supplementary Figure 1B). Notably, although no obvious SARS-CoV-2 replication in the pulmonary endothelium (Figure 2B), obvious cytokines release (including IL-7, CXCL10, CCL3 and IL-10) was still detected in the endothelial channel (Supplementary Figure 1B), mimicking the SARS-CoV-2-indcued cytokine storm *in vivo*. Compared with SARS-CoV-2 infection, H1N1 virus infection caused a milder increase of cytokines (Supplementary Figure 1B).

Then, the collected pulmonary endothelial medium was diluted with twice volume of fresh endothelial medium, and infused it into the vascular channel of BBB chips for continued culture. 4 days later, permeability assay showed infusion of endothelial medium from SARS-CoV-2-infected human alveolus chips induced a significant increase in BBB permeability, with a less increase in H1N1 virus group (Figure 3A).

**Figure 3.**
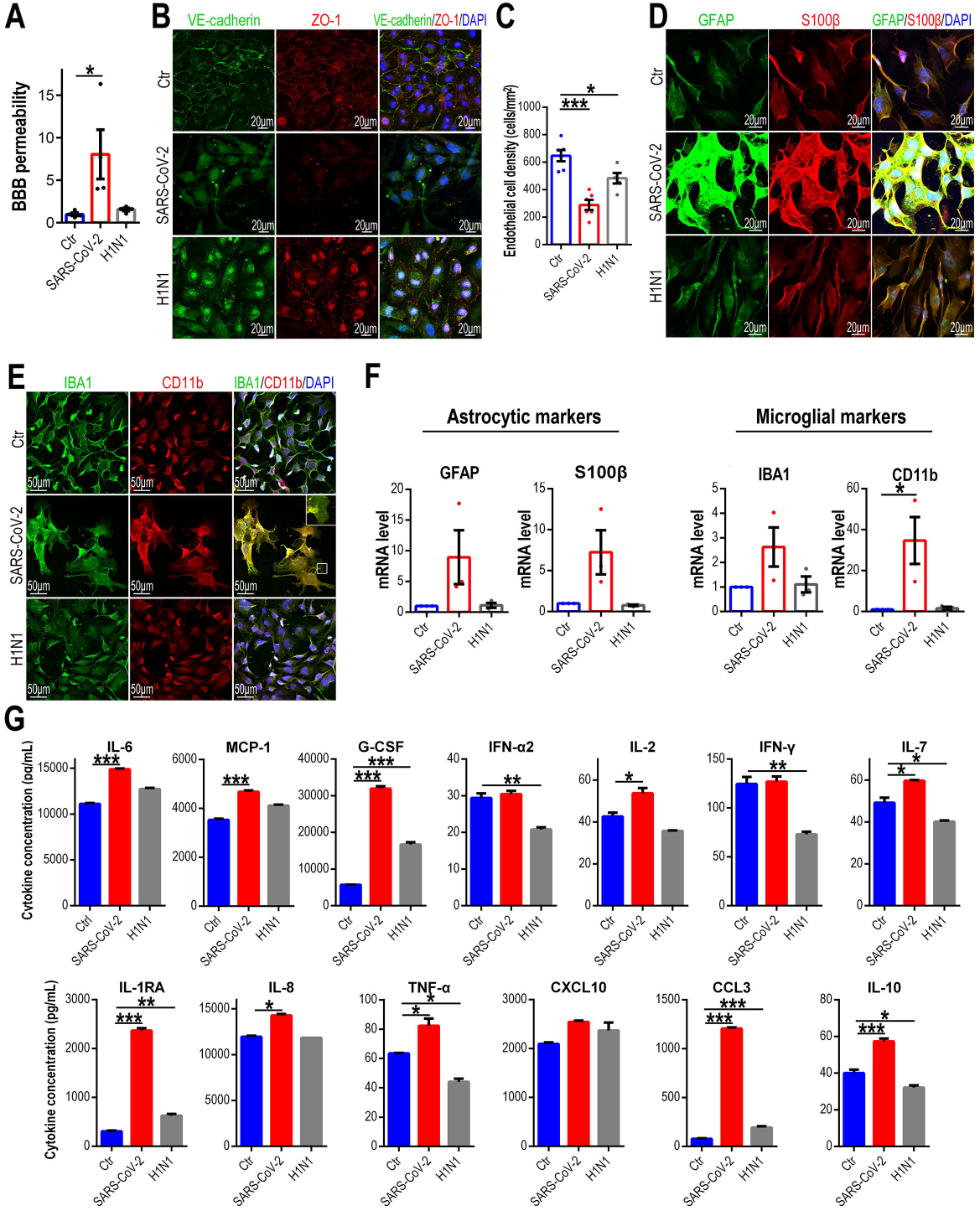
Exploring the indirect effect of SARS-CoV2 on BBB function after lung infection on the linked organ chip platform. (A) FITC-dextran permeability assays showed BBB permeability changes at Day 4 following infusion of endothelial medium from SARS-CoV-2 or H1N1 virus infected human alveolus chips (n=4). The FTIC-dextran concentration of lower channel in control group was normalized to 1, and that of SARS-CoV-2 or H1N1 virus group was presented as the multiples of FTIC-dextran concentration in the control group. Data are presented as mean±SEM and analyzed by unpaired Student’s t test (*: p<0.05). (B) Representative confocal micrographs of brain endothelial cells immunostained for VE-cadherin and ZO-1 at Day 4 following conditional medium treatment. (C) Quantification of the endothelial cell density based on (B). Data represent at least five independent experiments and more than 300 cells were quantified. Data are presented as mean±SEM and analyzed by unpaired Student’s t test (*: p<0.05; ***: p<0.001). (D) Representative confocal micrographs of astrocytes immunostained for GFAP and S100β at Day 4 following conditional medium treatment. (E) Representative confocal micrographs of microglia immunostained for CD11b and IBA1 at Day 4 following conditional medium treatment. The lamellipodia indicated by white box was enlarged in top right corner. (F) qRT-PCR results showing the mRNA level changes of GFAP, S100β, CD11b and IBA1 in glial cells at Day 4 following conditional medium treatment. Data are presented as mean±SEM (n=3) and analyzed using a one-way ANOVA with Bonferroni post hoc test (*: p<0.05). (G) Multiplex assays showed 13 cytokine levels in culture supernatants of CNS channel at Day 4 following conditional medium treatment (n=3). Data are presented as mean±SEM, and are analyzed using a one-way ANOVA with Bonferroni post-test (*: p<0.05; **: p<0.01; ***: p<0.001).

As brain microvascular endothelial cells are the core anatomical component of BBB, tight junctions and adherens junctions play critical roles in keeping adjacent endothelial cells close together and preventing paracellular diffusion. We then detected the tight junctions and adherens junction following conditional medium treatment, and the results showed endothelial tight junctions and adherens junctions were severely disrupted, as indicated by altered ZO-1 and VE-cadherin organization, with a lesser change in H1N1 virus group (Figure 3B). Also, the endothelial cell density showed an obvious decrease in SARS-CoV-2 group (Figure 3C).

### SARS-CoV-2 induced glial activation and neuroinflammation on the linked organ chip platform

To monitor the conditions of astrocytes and microglia following conditional medium treatment, glial cells were examined by staining with cell-specific markers. Confocal microscopic analysis showed, both astrocytic markers (GFAP, S100β) (Figure 3D) and microglial markers (IBA1 and CD11b) (Figure 3E) were significantly up-regulated in SARS-CoV-2 group, while without obvious changes in H1N1 virus group. Intriguingly, in SARS-CoV-2 group, we noticed many microglia were distributed in cluster, resembling the focal microglial activation *in vivo* characterized by microglial nodules ^14^. Besides, many lamellipodia (enlarged area in Figure 3E) were observed in the microglia of SARS-CoV-2 group, and similar phenomenon of activated microglia were also reported in previous studies ^45,46^. Consistently, the mRNA level of GFAP, S100β, IBA1 and CD11b were all specifically up-regulated in SARS-CoV-2 group by qRT-PCR analysis (Figure 3F). Besides, cytokine assay showed 10 cytokines (including IL-6, MCP-1, G-CSF, IL-2, IL-7, IL-1RA, IL-8, TNF-α, CCL3 and IL-10) were significantly released by the glial cells (CNS channel) in SARS-CoV-2 group (Figure 3G).

Since astrocytes and microglia were activated following pulmonary SARS-CoV-2 infection, we then investigated whether the activated glial cells were involved in the BBB damage, especially the brain endothelium disruption. Here, we ablated the astrocytes or microglia from the BBB chip, and examined their influences on brain endothelium following SARS-CoV-2 infection. As Figure 4A showed, 4 days after infusion of endothelial medium from SARS-CoV-2-infected alveolus chip, astrocytes or microglia ablation couldn’t restore the normal cell periphery distribution of VE-cadherin and ZO-1. However, astrocytes ablation partially ameliorated the brain endothelial cell detachment from the porous membrane, while microglia ablation showed a weaker rescue effect (Figure 4A, B). These results suggested, activated glial cells, especially activated astrocytes, exacerbated brain endothelial damage following SARS-CoV-2 infection on the linked chip platform.

**Figure 4.**
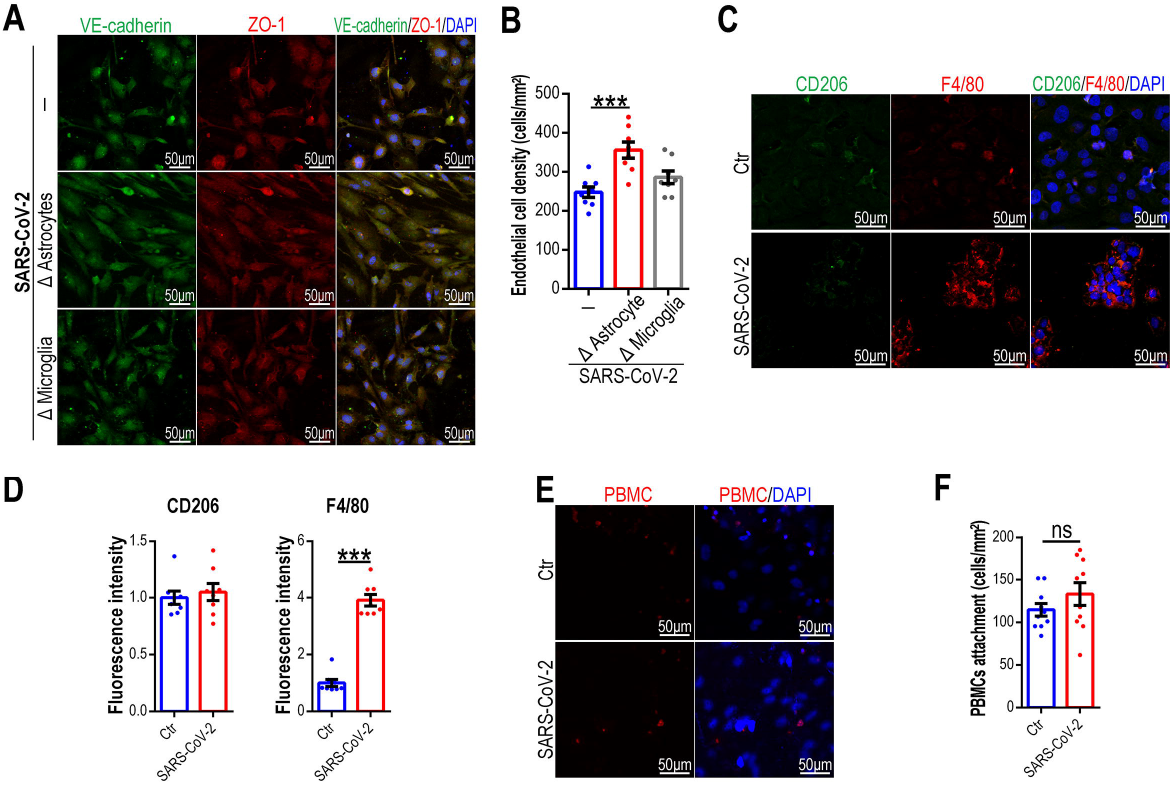
Characterizing immune responses of glial cells and immune cells on the linked organ chip platform. (A) Representative confocal micrographs of brain endothelial cells immunostained for VE-cadherin and ZO-1 without astrocytes (Δ Astrocytes) or without microglia (Δ Microglia) following infusion of endothelial medium from SARS-CoV-2-infected human alveolus chips at Day 4 (n=4). (B) Quantification of endothelial cell density based on (A). 8 randomly selected areas were quantified for each group. Data are presented as mean±SEM and analyzed using a one-way ANOVA with Bonferroni post hoc test (***: p<0.001). (C) Representative confocal micrographs of microglia immunostained for CD206 and F4/80 following infusion of endothelial medium from SARS-CoV-2-infected human alveolus chips at Day 4 (n=3). (D) Quantification of CD206 and F4/80 fluorescence intensity for control and SARS-CoV-2 group based on (C). 8 randomly selected areas were quantified for each group. Data are presented as mean±SEM and analyzed by unpaired Student’s t test (***: p<0.001). (E) Representative confocal micrographs of PBMCs (stained by Cell Tracker Red dye) attachment on endothelium following infusion of endothelial medium from SARS-CoV-2-infected human alveolus chips at Day 2 (n=3). (F) Quantification of density of attached PBMCs on endothelium based on (E). 10 randomly selected areas were quantified for each group. Data are presented as mean±SEM and analyzed by unpaired Student’s t test (ns: p>0.05).

As the resident immune cells of nervous system, microglia play vital roles in the maintenance of brain homeostasis, and responses to injury or infection. Upon activation, microglia can dynamically change their status into M1 phenotype (proinflammatory) or M2 phenotype (anti-inflammatory), a process termed as polarization ^47,48^. To define the status of microglia during BBB damage, we examined the marker expressions of M1 or M2 microglia by staining F4/80 and CD206, respectively. A significant increase in F4/80^+^ cells (M1 marker) were observed in SARS-CoV-2 group, with no obvious change for CD206^+^ cells (M2 marker) (Figure 4C, D), which indicated SARS-CoV-2 infection promoted M1 microglial polarization and induced neuroinflammation.

In addition to glial cells, peripheral immune cells may also participate in the BBB injury. To evaluate their roles, endothelial medium from SARS-CoV-2-infected alveolus chips containing ∼1×10^5^ PBMCs (pre-stained with Cell Tracker Red dye) were infused into the vascular channel of BBB chips, and 2 days later, the recruitment of peripheral immune was assessed by the PBMCs binding to the brain endothelium. The results showed, conditional medium treatment only leaded to a slight increase of PBMCs attachment on endothelium in the early stage of BBB injury (P value>0.05; Figure 4E, F).

All these findings indicated, SARS-CoV-2 could induce BBB injury and glial cell activation indirectly, which may be mediated by systematic inflammation after lung infection. Once the glial cells were activated, they exacerbated brain endothelial injury in turn and induced more severe inflammation.

## DISCUSSION

COVID-19 is a systematic disease with multi-organ symptoms, and neurological symptoms are common in a large population of patients. In this study, we proposed a new strategy to probe the effects of SARS-CoV-2 infection on human brains in a linked chip platform. This linked chip platform was established by combining a human alveolar chip with a human BBB chip, and enabled to recapitulate the basic features of alveolar-capillary barrier and blood-brain barrier *in vivo*. Our results revealed, direct SARS-CoV-2 exposure to vascular side had no visible effects on the BBB chip alone, while it can lead to disrupted BBB and neuroinflammation following lung infection indirectly on the linked chip platform. These new findings suggested the neurological alterations in COVID-19 may not be caused by direct viral neuroinvasion, but rather likely are the consequences of systematic inflammation following lung infection.

The pathological changes of central nervous system in COVID-19 patients have raised great concern. Many studies uncovered neurological changes by examining autopsy brain samples from COVID-19 patients, and some of our findings are consistent with these pathological reports. A study reported, cerebral injuries, such as hypoxic/ischemic damage, hemorrhage and glial activation, can be detected in COVID-19 brain samples, whereas RNAscope and immunocytochemistry failed to detect SARS-CoV-2 ^14^. Even though qRT-PCR examination showed low or trace levels of viral RNA in some COVID-19 brains, the levels of detectable virus were not associated with the severity of neuropathological alterations ^14,15^. Similarly, a recent study of single-nucleus transcriptomes revealed no molecular traces of virus in the frontal cortex and choroid plexus samples of COVID-19 patients either ^49^. All these findings indicated, the direct viral neuroinvasion may not be the prerequisite for the neurological manifestations of COVID-19. Notably, some pathological responses detected in our organ chip model, such as activated astrocytes with high degree of GFAP expression and activated microglia distributed in microglial nodules, were reminiscent of the similar phenomenon in brain tissues of COVID-19 patients ^14,15^, which verified the feasibility of our model in recapitulating neurological alterations in patients.

In human, BBB functions as a protective interface, prevents toxins and pathogens (including virus) from accessing the brain parenchyma. In regard to the viral neuroinvasion, several organoid models have been utilized to investigate the SARS-CoV-2 neurotropism and viral entry route to human brains ^19-24^, however, some limitations constrain interpretation of these studies. As these brain organoids have not realized vascularization and lack a functional BBB interface, making it difficult to accurately recapitulate the viral exposure in physiological state. Besides, no immune cells (such as peripheral blood immune cells, microglial cells) were present in these brain organoids, which made it unsuitable to evaluate the host immune responses and the impacts of systematic inflammation on brain following SARS-CoV-2 infection.

Compared with other *in vitro* models, the major advantage of this platform is the capability to introduce various of cell types or tissues according to research needs, such as immune cells, tissue interfaces and vasculatures. As for COVID-19, although immune cells and vascular endothelial cells are not the primary attack targets for virus, they contribute a lot to disease progression and some lifethreatening complications, such as inflammatory responses, systemic vasculitis and venous thromboembolism et al. ^50-52^. In this model system, peripheral blood immune cells were introduced into the alveolus chips, and robust inflammatory cytokines release was detected following SARS-CoV-2 infection, mimicking the cytokine storms commonly happening in severe COVID-19 cases. In our previous study on human alveolus chip model alone, the inclusion of peripheral immune cells significantly exacerbated the cytokines release and SARS-CoV-2-induced alveolar injury ^38^. Regarding vascular endothelial cells, our studies revealed exposure to the pulmonary endothelial medium containing pro-inflammatory cytokines triggered a second release of cytokines and chemokines in the vascular channel of BBB chips (Supplementary Figure 2). Through this amplification loop of inflammatory response, the brain endothelium constituted a significant source of inflammatory cytokines in human brains, such as IL-1, IL-6 and TNF-α that characterize the cytokine storm in COVID-19. All these findings indicated immune cells and endothelial cells are two major characters involved in COVID-19 pathogenesis.

It’s worth noting that, in the study we only discussed one potential entry route of SARS-CoV-2 to human brains actually. In addition to BBB, there is another barrier system in human central nervous system: blood-cerebrospinal fluid barrier (B-CSF-B), which separates the stroma capillaries from the CSF ^23,53^. Compared with the highly selective and insulated BBB, the B-CSF-B is much simpler, and formed by a single epithelial layer of the choroid plexus ^53,54^. Two groups reported, SARS-CoV-2 can infect choroid plexus epithelial cells and invade human brains by disrupting B-CSF-B ^23,24^. Some pathological studies also reported viral RNA can be detected in the cerebrospinal fluid or brains of some COVID-19 patients ^8,55,56^. However, the presence of virus in human brains or CSF is not a widely reported finding, especially compared with the huge population of COVID-19 patients suffering neurological symptoms. Based on the studies using the linked organ chip platform, we proposed that neuroinvasion of SARS-CoV-2 is not the prerequisite of neuropathological alteration in COVID-19 patients, and systematic inflammation is sufficient to induce BBB disruption and neuroinflammation in humans. In the course of disease progression, BBB sensed and relayed peripheral inflammation into the brain parenchyma, and the activated glial cells exacerbated the neuroinflammation.

Collectively, this work provided a proof of concept to utilize a linked organ chip model to study the influences of SARS-CoV-2 infection on human brains in an organ-organ context. This *in vitro* model system can not only simulate physiologically relevant structures and functions of biological barriers in a biomimetic microfluidic condition, but also recapitulate the complex pathophysiological changes following SARS-CoV-2 infection. We envision this microengineered linked organ model can provide a unique platform for the study of host-virus interactions, infection mechanisms and drug testing in particularly for virology research.

## METHODS AND MATERIALS

### Cell culture

Vero E6 cells were obtained from American Type Culture Collection (ATCC; no. 1586), and maintained in MEM medium (Gibco) supplemented with 10% fetal bovine serum (FBS; Gibco) and 1% Penicillin/Streptomycin (P/S). HPAEpiC were purchased form ScienCell Corporation (no. 3200), and maintained in RPMI 1640 medium (Gibco) supplemented with 10% FBS and 1% P/S. HULEC-5a cells were purchased from Procell Corporation (no. CL-0565; introduced from ATCC Company), and maintained in a special pulmonary endothelial medium (Procell Corporation, no. CM-0565). HBMEC cells were purchased from ScienCell Corporation (no. 1000), and maintained in ECM medium (ScienCell; no. 1001) supplemented with 10% FBS, 1% ECGS and 1% P/S. HA cells were a kind gift from Dr. Dan Liu’s lab (Huaqiao University), which was purchased from ScienCell Corporation (no. 1800), and maintained in AM medium (ScienCell; no. 1801) supplemented with 2% FBS, 1% AGS and 1% P/S. HMC3 cells were purchased from Procell Corporation (no. CL-0620), and maintained in MEM medium containing NEAA (Procell; no. MP150410) supplemented with 10% FBS and 1% P/S. Human peripheral blood mononuclear cells (PBMCs) were isolated from fresh human peripheral blood using Ficoll (GE Healthcare) density gradient centrifugation, and were cultured in RPMI 1640 medium containing 10% FBS, 1% P/S and 50 IU/mL IL-2. All cells were cultured at 37 °C in a humidified atmosphere of 5% CO2.

### Device fabrication

The human BBB chip device consists of an upper and lower layer fabricated by casting Polydimethylsiloxane (PDMS) pre-polymer on molds prepared using conventional soft lithography procedures. 10:1 (wt/wt) PDMS base to curing agent (184 Silicone Elastomer, Dow Corning Corp) was polymerized to produce molded device with channels by thermal curing at 80 °C for 45 min. The top channel (1.5 mm wide × 0.2 mm high) and bottom channel (1.5 mm wide × 0.2 mm high) are used to form the vascular lumen and the CNS layer, respectively. The overlapping area of upper and lower channels is 26.2985 mm^2^. The two channels are separated by a thin (10 μm thick) and porous (2 μm diameter) PET membrane to construct the tissue-tissue interface. Before seeding cells on the chips, the chips were sterilized by ultraviolet irradiation overnight and then were pre-coated with 50 μg/mL fibronectin at 37 °C for 24 hours.

### Virus

For SARS-CoV-2 infection experiments, a clinical isolate strain 107, which was obtained from Guangdong Provincial Center for Disease Control and Prevention, Guangdong Province of China. The virus was propagated in Vero E6 cells, and viral titer was determined by a TCID50 assay on Vero E6 cells. All the SARS-CoV-2 infection experiments were performed in the biosafety level-3 (BSL-3) laboratory.

For H1N1 virus infection experiments, the influenza A virus (IAV: H1N1 PR8) was provided by the National Institute for Viral Disease Control and Prevention (China). All the H1N1 virus infection experiments were performed in the BSL-2 laboratory.

### Cell culture on organ chips

To construct the human alveolus chip, HULEC-5a cells (∼5×10^4^ cells/chip) were initially seeded on the bottom side of the fibronectin-coated porous PET membrane and allowed to attach on the membrane surface under static conditions. Two hours later, HPAEpiC cells (∼1×10^5^ cells/chip) were seeded into the upper channel under static cultures. After cell attachment, constant media flows (100 μl/h) were applied in both the upper and bottom layers using a LongerPump. The cells were cultured for 3 days to form confluent alveolar-capillary barrier interface.

To construct the human BBB chip, HA cells (∼5×10^4^ cells/chip) were initially seeded on the bottom side of the fibronectin-coated porous PET membrane and allowed to attach on the membrane surface under static conditions. Two hours later, the lower channel was washed with fresh medium to remove the unattached HA cells. Then, HMC3 cells (∼5×10^4^ cells/chip) were seeded on the bottom side of lower channel, and HBMEC cells (∼1×10^5^ cells/chip) were seeded into the upper channel, respectively. After cell attachment, constant media flows (100 μ l/h) were applied in both the upper and bottom layers using a LongerPump. The cells were cultured for 3 days to form confluent blood-brain barrier interface.

All the chips were maintained in an incubator with 5% CO_2_ at 37°C.

### Virus infection on chips

For virus infection on human alveolus chips, 20 μL RPMI 1640 medium containing the indicated multiplicity of virus (SARS-CoV-2 or H1N1 virus) were infused in the upper channel. One hour after infection, cells in the upper channel were washed two times with PBS and kept in fresh medium for continued culture.

For virus infection on human BBB chips, 20 μL ECM medium containing the indicated multiplicity of virus (SARS-CoV-2) were infused in the upper channel. One hour after infection, cells in the upper channel were washed two times with PBS and kept in fresh medium for continued culture.

### Endothelial medium collection from infected human alveolus chips

Firstly, 100 μL fresh pulmonary endothelial medium containing ∼1×10^5^ PBMCs were infused in the lower vascular channel. Then, SARS-CoV-2 or H1N1 virus (MOI=1) was inoculated in the upper alveolar channels. One hours later, the upper alveolar channels were washed twice with PBS, and then 100 μL fresh RPMI 1640 medium was infused into the upper alveolar channels. Every two days (Day 2, 4, 6 post-infection), endothelial medium (conditional medium) was collected, centrifuged (10,000 g; 10 min) and stored as at -80 °C. 100 μL fresh pulmonary endothelial medium containing ∼1×10^5^ PBMCs and 100 μL fresh RPMI 1640 medium was infused into the lower and upper channels respectively for continued culture.

### Infusion of endothelial medium from the infected alveolus chips into the BBB chips

Pulmonary endothelial culture supernatants from the infected alveolus chips (Mock, SARS-CoV-2 or H1N1 virus-infected) were diluted with twice volume of fresh ECM medium. Every two days, 100 μL diluted conditional medium and 100 μL fresh glial cell medium (HA cell medium : HMC3 cell medium=1:1) was infused into the upper and lower channels of BBB chips, respectively for continued culture (4 days).

### Immunofluorescent imaging of organ chips

Cells were washed twice with PBS through the upper and lower channels and fixed with 4% PFA overnight. The fixed cells were permeabilized and blocked with PBST (0.2% Triton X-100 in PBS) buffer containing 5% normal goat serum for 30 min at room temperature. Antibodies were diluted with PBST buffer and infused into the upper and lower channels, respectively. Cells were stained with the primary antibodies at 4 °C overnight, and with the secondary antibodies at room temperature for 1 h. After counterstained with DAPI, chips were disassembled, and porous membrane and lower channel were mounted on slides respectively. All images were acquired using a Carl Zeiss LSM880 confocal fluorescent microscope system. Image processing was done using ImageJ software (NIH).

### Viral RNA copies detection by qRT-PCR

100 μL culture supernatants were harvested separately from the upper and lower channels of chips, and extracted for viral RNA using the HP Viral RNA Kit (Roche, Cat no. 11858882001) according to the manufacturer’s instructions. qRT-PCR was performed on Real-Time PCR System (Applied Biosystem, ViiA™ 7) with One Step RT-PCR RNA direct realtime PCR mastermix (TOYOBO, QRT-101A). The primers and probe for qRT-PCR were used as follows: N-F: 5’-GGGGAACTTCTCCTGCTAGAAT-3’; N-R: 5’-CAGACATTTTGCTCTCAAGCTG-3’; SARS-CoV-2 N-probe: 5’-TTGCTGCTGCTTGACAGATT-3’. PCR amplification was performed as follows: 50°C for 10 min, and 95°C for 1min followed by 45 cycles consisting of 95°C for 15 s, 60°C for 45 s.

### Permeability assay

The BBB chip permeability was assessed by detecting the diffusion of 40 kDa FITC-dextran from the upper vascular layer to the lower parenchyma channel. Briefly, ECM medium containing 500 μg/mL FITC-dextran was infused into the upper vascular channel of BBB chip. 2 hours later, the media of lower channel was collected and the FITC-dextran concentration was determined by FITC fluorescence intensity using a microplate reader system (BioTek Synergy H1) at 490 nm (Ex) and 525 nm (Em).

### Multiplex assay for cytokines detection

Culture supernatants were harvested from upper channels and lower channels of chips, separately. Cytokines in culture supernatants, including IL-6, MCP-1 (CCL2), G-CSF, IFN-α2, IL-2, IFN-γ, IL-7, IL-1RA, IL-8 (CXCL8), TNF-α, CXCL10 (IP-10), CCL3 (MIP-1α) and IL-10 were analyzed using a human COVID-19 Cytokine Storm Panel 1 (13-plex) kit (Cat. No. 741091; BioLegend, USA) according to the manufacturer’s instructions.

### Western blot

Protein samples were separated on 10% SDS-PAGE and then transferred onto 0.2 μm nitrocellulose membranes (GE Amersham). After being blocked with 5% non-fat powdered milk in TBST buffer containing 0.05% Tween-20, the membranes were probed with the primary antibodies at 4 °C overnight. The membranes were then probed with corresponding horseradish peroxidase (HRP)-conjugated secondary antibodies at room temperature for 1 h at room temperature. Protein bands were detected by Prime Western Blotting Detection Reagent (GE life).

### Statistical analyses

Data were collected by Excel (Microsoft) software. The GraphPad Prism 6 software was used for data statistical analysis. All experiments were performed at least three times. During the experiment and assessing the outcome, the investigators were blinded to the group allocation. Differences between two groups were analyzed using unpaired Student’s t test. Multiple group comparisons were performed using a one-way analysis of variance (ANOVA) followed by Bonferroni post hoc test. The bar graphs with error bars represent mean ± standard error of the mean (SEM). Significance is indicated by asterisks: *, P < 0.05; **, P < 0.01; ***, P < 0.001.

## ACKNOWLEDGEMENT

This research was supported by the Strategic Priority Research Program of the Chinese Academy of Sciences,Grant (Nos. XDB32030200, XDB29050301, XDA16020900), National Key R&D Program of China (No. 2017YFB0405404), Yunnan Key Research and Development Program (No.202003AD150009), National Nature Science Foundation of China (No. 3210100629, 31971373), China Postdoctoral Science Foundation (No.2019M660065), Innovation Program of Science and Research from the DICP, CAS (DICP I201934).

## CONFLICT OF INTEREST

The authors declare that they have no conflict of interest.

## Notes

### Competing Interest Statement

The authors have declared no competing interest.

