## Supplemental File for "SARS-CoV-2 causes human BBB injury and neuroinflammation indirectly in a linked organ chip platform"

^2^Key Laboratory of Animal Models and Human Disease Mechanisms of Chinese Academy of Sciences/Key Laboratory of Bioactive Peptides of Yunnan Province, KIZ-CUHK Joint Laboratory of Bioresources and Molecular Research in Common Diseases, National Resource Center for Non-Human Primates, Kunming Primate Research Center, National Research Facility for Phenotypic & Genetic Analysis of Model Animals (Primate Facility), Sino-African Joint Research Center, and Engineering Laboratory of Peptides, Kunming Institute of Zoology, Chinese Academy of Sciences, Kunming 650107, Yunnan, China

^3^Kunming National High-level Biosafety Research Center for Non-Human Primates, Kunming Institute of Zoology, Chinese Academic of Sciences, Kunming, Yunnan 650107, China

^4^University of Chinese Academy of Sciences, Beijing, China

^5^Kunming National High-level Bio-safety Research Center for Non-human Primates, Center for Biosafety Mega-Science, Kunming Institute of Zoology, Chinese Academy of Sciences, Kunming, 650107, China

^6^Core Technology Facility of Kunming Institute of Zoology, Chinese Academy of Sciences，Kunming 650223, Yunnan, China

^7^Institute for Stem Cell and Regeneration, Chinese Academy of Sciences, Beijing, China

^8^CAS Center for Excellence in Brain Science and Intelligence Technology, Chinese Academy of Sciences, Shanghai, China

***Correspondence:**

Jianhua Qin, Division of Biotechnology, Dalian Institute of Chemical Physics, Chinese Academy of Sciences, 457 Zhongshan Road, Dalian 116023, China.. Tel: 86-0411-84379650.

Ren Lai, Kunming Institute of Zoology, the Chinese Academy of Sciences, No. 32 Jiaochang Donglu, Kunming, 650223, China.. Tel: 86-0871-65197578.

^#^These authors contributed equally.

**Supplementary Table 1. Primer sequences used for RT-qPCR analysis in this study.**

| **Primer** | **Sequence (5’-3’)** |
| --- | --- |
| GFAP-F | ACCTGCAGATTCGAGAAACC |
| GFAP-R | CTCCTTAATGACCTCTCCATCC |
| S100β-F | GAGGAATGAAGGGCCACTGA |
| S100β-R | CCCTAGGCACCAGCAGGTC |
| IBA1-F | GATGATGCTGGGCAAGAGAT |
| IBA1-R | CCTTCAAATCAGGGCAACTC |
| CD11b-F | TGAAGGCTCAGACAGAGACCAAA |
| CD11b-R | GCTGCCCACAATGAGTGGTAC |
| GAPDH-F | CTTCACGACCATGGAGAAGG |
| GAPDH-R | CCAAGCAGTTGGTGGTGCAG |

**Supplementary Table 2. Primary antibodies used for immunofluorescence in this study.**

| **Antibody** | **Vendor** | **Catalog #** | **Dilution** |
| --- | --- | --- | --- |
| Anti-ZO-1 | Abcam | Ab96587 | 1:100 |
| Anti-VE-cadherin | Proteintech Group | 66804-1-Ig | 1:100 |
| Anti-VE-cadherin | Cell Signaling Technology | 2500S | 1:100 |
| Anti-GFAP | Cell Signaling Technology | 3670S | 1:100 |
| Anti-GFAP | Proteintech Group | 16825-1-AP | 1:100 |
| Anti-S100β | Proteintech Group | 15146-1-AP | 1:100 |
| Anti-IBA1 | Abcam | Ab178847 | 1:100 |
| Anti-IBA1 | Proteintech Group | 66827-1-Ig | 1:100 |
| Anti-CD11b | Abcam | ab52478 | 1:100 |
| Anti-E-cadherin | Proteintech Group | 60335-1-Ig | 1:100 |
| Anti-Spike | Sino Biological | 40150-R007 | 1:200 |
| Anti-H1N1 M2 | GeneTex | GTX125951 | 1:1000 |
| Anti-F4/80 | Proteintech Group | 28463-1-AP | 1:100 |
| Anti-CD206 | Proteintech Group | 60143-1-Ig | 1:100 |

**Supplementary Table 3. Primary antibodies used for Western blot in this project.**

| **Antibody** | **Vendor** | **Catalog #** | **Dilution** |
| --- | --- | --- | --- |
| Anti-ACE2 | Proteintech Group | 21115-1-AP | 1:1000 |
| Anti-TMPRSS2 | Proteintech Group | 14437-1-AP | 1:1000 |
| Anti-GAPDH | CWBIO | CW0100 | 1:2000 |

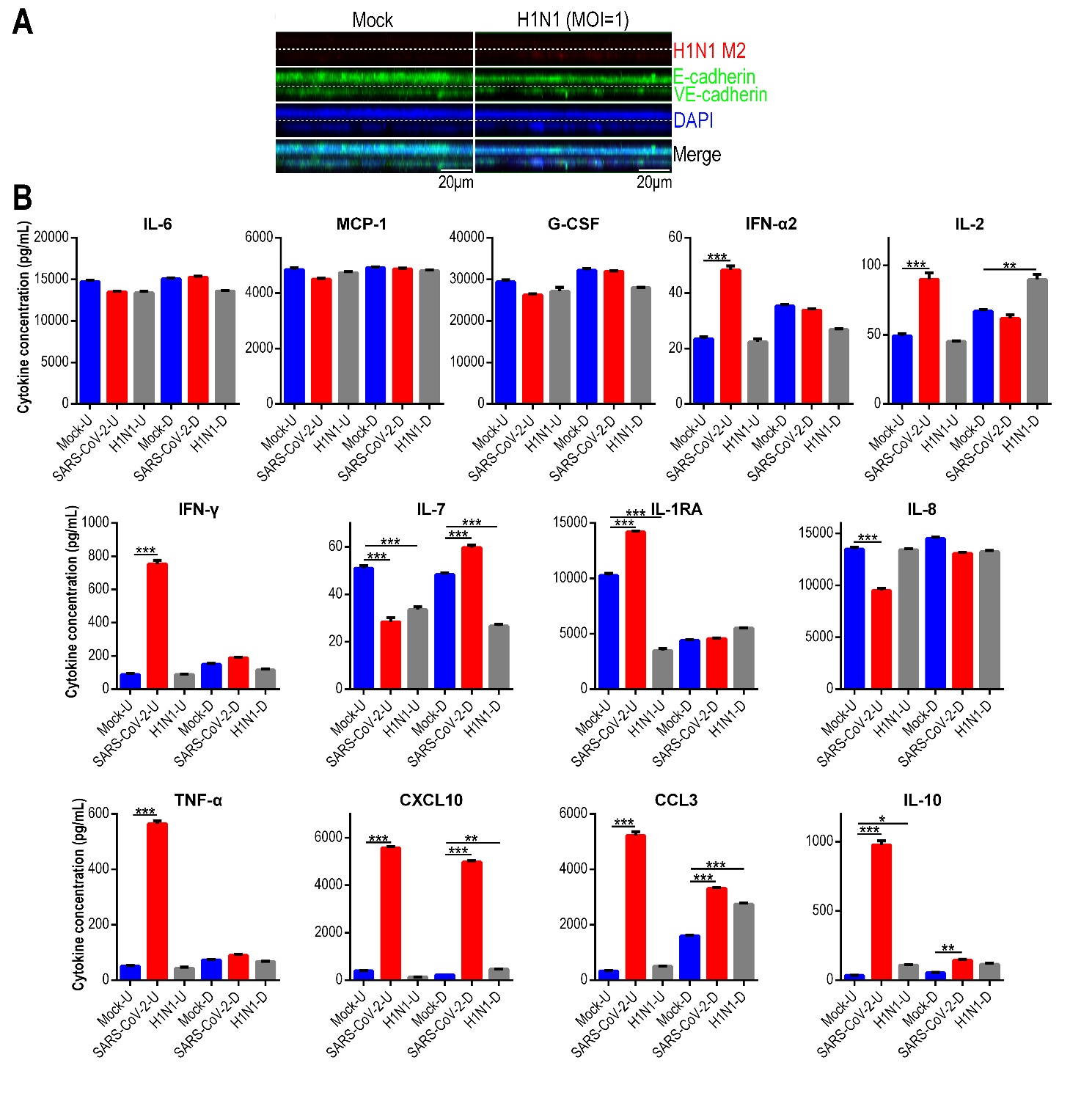

**Supplementary Figure 1. SARS-CoV-2 infection or H1N1 infection on human alveolus chip.** (A) Representative side view of the alveolar-capillary barrier 3 days after H1N1 virus (PR8 strain) infection (MOI=1) on alveolus chip. (B) Multiplex assays showed 13 cytokine levels in culture supernatants of upper epithelial channel (-U) and lower endothelial channel (-D) for mock- and SARS-CoV-2-infected BBB chips at Day 4 post-infection (n=3). Data are presented as mean±SEM, and are analyzed using a one-way ANOVA with Bonferroni post-test (*: p<0.05; **: p<0.01; ***: p<0.001).

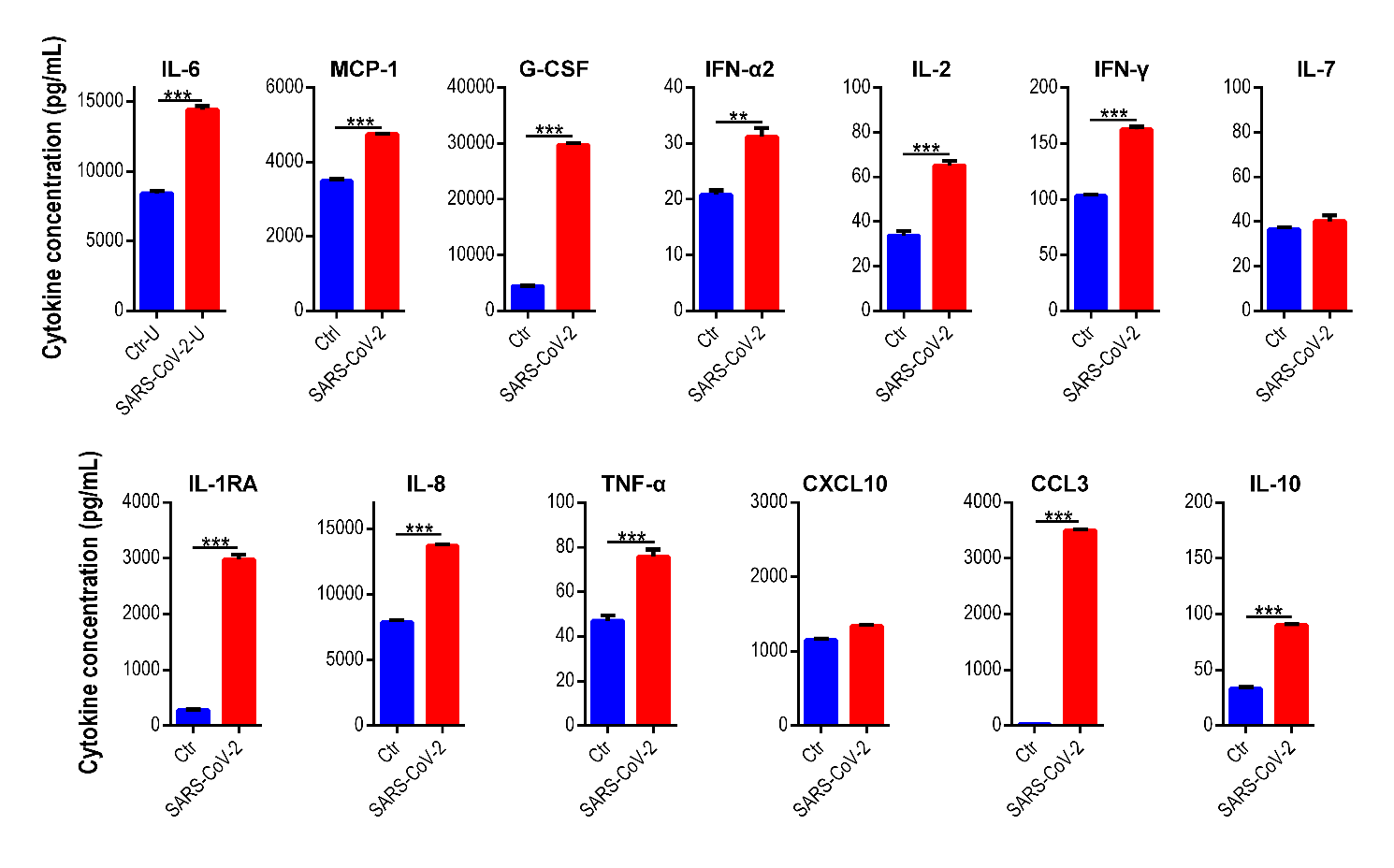

**Supplementary Figure 2. Inflammatory cytokines release in vascular channel of BBB chip following infusion of endothelial medium from SARS-CoV-2-infected alveolus chip.** Multiplex assays showed 13 cytokine levels in culture supernatants of vascular channel in BBB chips at Day 4 following conditional medium treatment (n=3). Data are presented as mean±SEM, and are analyzed using a one-way ANOVA with Bonferroni post-test (*: p<0.05; **: p<0.01; ***: p<0.001).
